# The effect of circulating neutralizing antibodies on the replication of SARS-CoV-2 variants following post-vaccination infections

**DOI:** 10.1101/2024.06.18.599357

**Authors:** Miguel A. Garcia-Knight, J. Daniel Kelly, Scott Lu, Michel Tassetto, Sarah A. Goldberg, Amethyst Zhang, Jesus Pineda-Ramirez, Khamal Anglin, Michelle C. Davidson, Jessica Y. Chen, Maya Fortes-Cobby, Sara Park, Ana Martinez, Matthew So, Aidan Donovan, Badri Viswanathan, Eugene T. Richardson, David R. McIlwain, Brice Gaudilliere, Rachel L. Rutishauser, Ahmed Chenna, Christos Petropoulos, Terri Wrin, Steve G. Deeks, Glen R. Abedi, Sharon Saydah, Jeffrey N. Martin, Melissa Briggs-Hagen, Claire M. Midgley, Michael J. Peluso, Raul Andino

**Author notes:** **Corresponding author contact information:** Raul Andino, University of California, San Francisco, MBGH S572E, Box 2280, 600 16^th^ Street, San Francisco, CA 94158, USA. **Conflict of interest statement:** The authors have declared that no conflict of interest exists.

## Abstract

The impact of pre-existing neutralizing antibodies (NAbs) titers on SARS-CoV-2 viral shedding dynamics in post-vaccination infection (PVI) are not well understood. We characterized viral shedding longitudinally in nasal specimens in relation to baseline (pre/peri-infection) serum neutralizing antibody titers in 125 participants infected with distinct SARS-CoV-2 variants. Among 68 participants who had received vaccinations, we quantified the effect of baseline serum NAb titers on maximum viral RNA titers and on the duration of infectivity. Baseline NAb titers were higher and efficiently targeted a broader range of variants in participants who received one or two monovalent ancestral booster vaccinations compared to those with a full primary vaccine series. In participants with Delta variant infections, baseline NAb titers targeting Delta were negatively correlated with maximum viral RNA copies. Per log_10_ increase in baseline NAb IC50, maximum viral load was reduced -2.43 (95% confidence interval [CI] -3.76, -1.11) log10 N copies and days of infectious viral shedding were reduced -2.79 [95% CI: -4.99, -0.60] days. By contrast, in those with Omicron infections (BA.1, BA.2, BA.4 or BA.5 lineages) baseline NAb responses against Omicron lineages did not predict viral outcomes. Our results provide robust estimates of the effect of baseline NAbs on the magnitude and duration of nasal viral replication after PVI (albeit with an unclear effect on transmission) and show how immune escape variants efficiently evade these modulating effects.

## INTRODUCTION

Circulating neutralizing antibodies (NAbs) against SARS-CoV-2 are associated with protection against infection and disease and are induced following both SARS-CoV-2 infections and vaccination^1^. However, NAbs titers wane in the months following their induction (through vaccination or infection) and time since vaccination correlates negatively with protection^2,3^. In addition, variants such as those descending from the B.1.1.529 (Omicron) lineage (with >30 amino acid mutations in the Spike protein relative to Wuhan-Hu-1) can evade NAbs targeting ancestral lineages and vaccine antigens^4^. This has contributed to widespread post-vaccination infections (PVI), reinfections and ongoing waves of community transmission, despite the use of distinct vaccine platforms and the provision of booster vaccinations with updated vaccine antigens^5^. However, most vaccine doses received globally, either as part of a primary vaccine series or through booster doses, have used antigens derived from ancestral Wuhan-Hu-1^6^.

Numerous studies have assessed the effect of vaccination on viral replication dynamics and infectiousness - key parameters linked to SARS-CoV-2 pathogenesis and transmission. However, few studies have been able to directly assess how these outcomes relate to the NAb response. For instance, the mRNA vaccine BNT162b2 can impact peak viral load in PVI early after vaccination^7^, though the effect is transient and not observed in all studies of outpatient cohorts^8,9^. Likewise, data on the effect of mRNA vaccines on the duration of viral shedding and infectiousness following PVI are conflicting^10^, with no reduction in the incidence of household transmission in Delta infections^9,11^ particularly 12 weeks after vaccination^12^. In the present study, we determine the relationship between NAb titers -elicited following vaccination and measured at the time of infection- and key virological parameters in a longitudinal household cohort sampled intensely during the period of viraemia. Our findings help quantify the protective effect of circulating NAbs against SARS-CoV-2 variants with implications for the study of COVID-19 pathogenesis and for efforts to model SARS-CoV-2 transmission in the context of novel prophylactic and therapeutic interventions.

## RESULTS

### Cohort characteristics

A total of 174 participants (125 SARS-CoV-2-infected and 49 uninfected) from 78 households were enrolled and had blood collected from September 2020 through September 2022 in the San Francisco Bay Area. A median of 13 (range 3-15) nasal specimens and 4 (range 1-4) blood specimens were collected per participant, totaling 2471 and 664 specimens, respectively. The demographic characteristics and vaccination histories of infected and uninfected participants are shown in **Supp. table 1**. Participants were infected with distinct variants including those that predated the emergence of variants of concern (preVOC), Epsilon, Delta and Omicron (sub-lineages BA.1, BA,2, BA.4 or BA.5). Among infected participants, 37/125 (29.6%) received a primary vaccine series and 31/125 (24.8%) received one or two original monovalent booster doses. A total of 45/57 (85%) unvaccinated participants were infected with preVOC viral lineages, 32/37 (87%) of those who received a primary vaccine series were infected with Delta variants and 31/31 (100%) participants who received original monovalent booster vaccinations had infections with Omicron variants.

### Viral shedding kinetics over the acute infection period

Among infected participants, we quantified viral RNA and assessed the presence of infectious virus longitudinally, by vaccine status (**Fig.1A**). Maximum viral RNA copies and the duration of infectious virus shedding did not differ significantly by vaccine status (**Fig. 1B & C**). Analysis of kinetics by variant, indicated that participants with BA.1 infections had a significantly reduced maximum RNA load than those with pre-BA.1 infections or those with BA.2/BA.4 and BA.5 infections (**Fig. 1D**). No differences were observed between variants with regards to the duration of infectious virus shedding (**Fig. 1E**).

**Figure 1.**
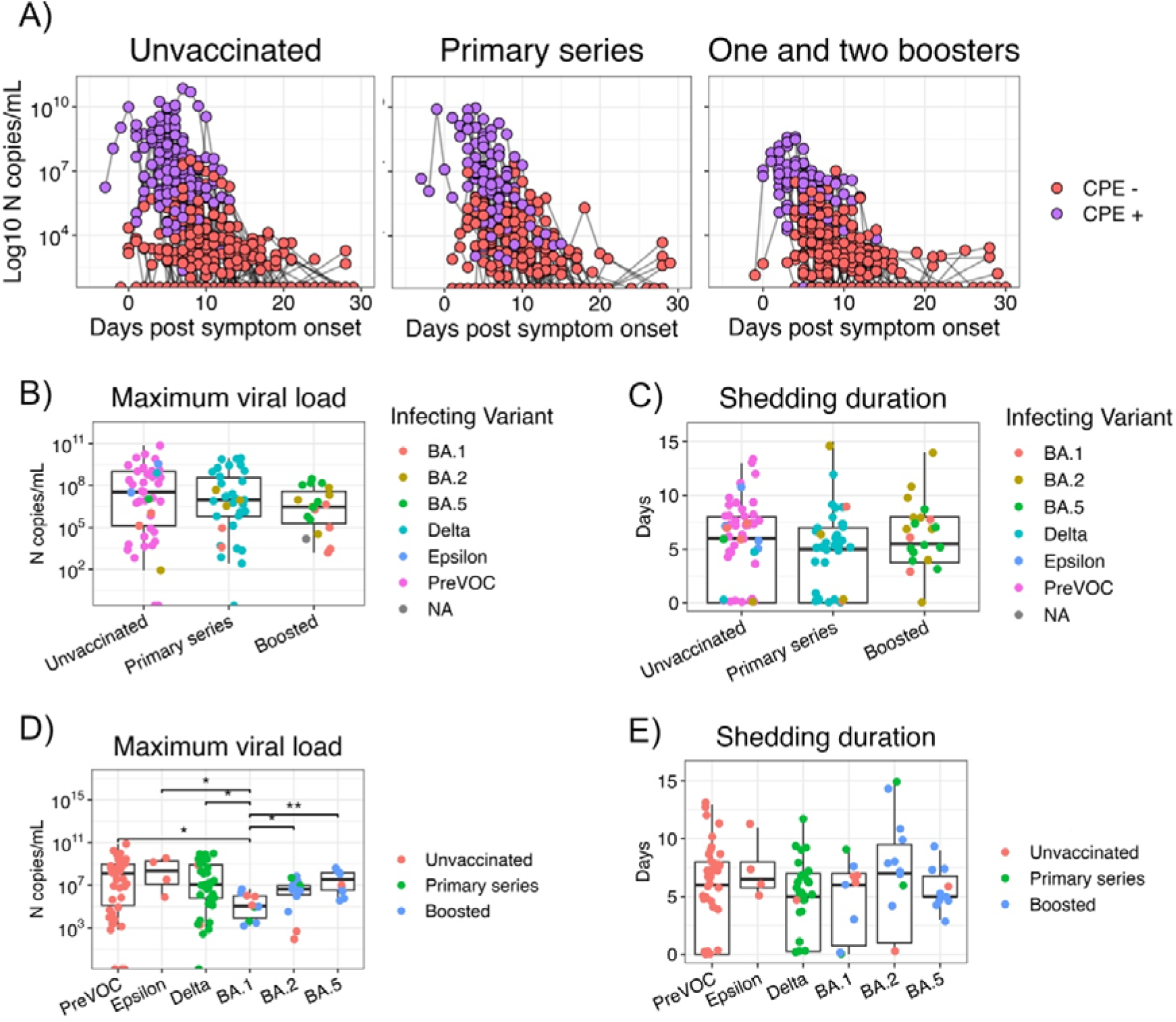
The influence of vaccination history and viral variant on viral shedding dynamics. A) Longitudinal viral shedding dynamics in nasal specimens collected over 28 days from symptom onset in unvaccinated participants (N=57) and participants with post-vaccination infections (PVI) who received a primary vaccine series (N=37) or monovalent booster vaccinations (N=31). Copies of SARS-CoV-2 nucleocapsid (N) RNA of each specimen and the presence of infectious virus are shown. B) Comparison of median maximum copies of N RNA between vaccine groups. C) Comparison of the median duration in days which infectious virus was detected between vaccine groups. D) Comparison of median maximum copies of N RNA in participants stratified by infecting variant. Vaccination history is indicated by the colour shown in the legend. E) Comparison of median duration in days which infectious virus was detected stratified by infecting variant. Vaccination history is indicated the colour shown in the legend. All pairwise comparisons were made using a two-sided Wilcoxon rank sum test. Only statistically significant differences are shown. *P < 0.05 and **P < 0.01. CPE, cytopathic effect.

### Baseline neutralizing antibody titers in participants with post-vaccine infections

Recruitment specimens were collected a median of 5 (interquartile range [IQR]: 4-5) days post symptom onset (PSO) in infected individuals. To focus on NAb titers prior to the induction of the anamnestic response following PVI, we excluded from our analysis participants (N=22) with recruitment specimens obtained on day 7 PSO or later^13^. No significant differences were observed in median NAb titers in recruitment specimens obtained <7 days PSO between participants with PVI and uninfected participants stratified by vaccination status (**Supp. Fig. 1**), indicating that baseline titers <7 days post-onset in infected participants resembled those prior to infection rather than reflecting post-infection responses. In addition, there was no difference in the time since last vaccine dose between PVI groups (**Supp. Fig 2**)

We initially assessed the strength and breadth of the baseline NAb response (**Fig.2A**). We observed that in participants with a primary vaccine series, titers against Omicron BA.1 and BA.2 were reduced 6.6 (P>0.0001) and 3.2 (P<0.001) fold, respectively, compared to Wuhan-Hu-1, and tended to be reduced 2.0 (P=0.08) fold against Beta. By contrast, in participants who received booster vaccinations, no reduction in baseline NAb titers was observed against Beta, and NAb titers were only reduced 2.4 (P=0.005) and 1.8 (P=0.001) fold against Omicron BA.1 and BA.2, respectively (**Fig.2A**). Overall, 17/37 (46%) and 10/37 (27%) participants with a primary vaccine series had undetectable NAb responses to BA.1 and BA.2, respectively, compared to 1/31 (3%) and 0/31 (0%) of participants who received booster vaccinations. Baseline NAb titers targeting the infecting variant for each participant (except those with BA.5 infections for which response against BA.2 are shown) were higher in those that had received vaccine booster doses compared to those who received a primary series (P<0.05) (**Fig.2B**).

**Figure 2.**
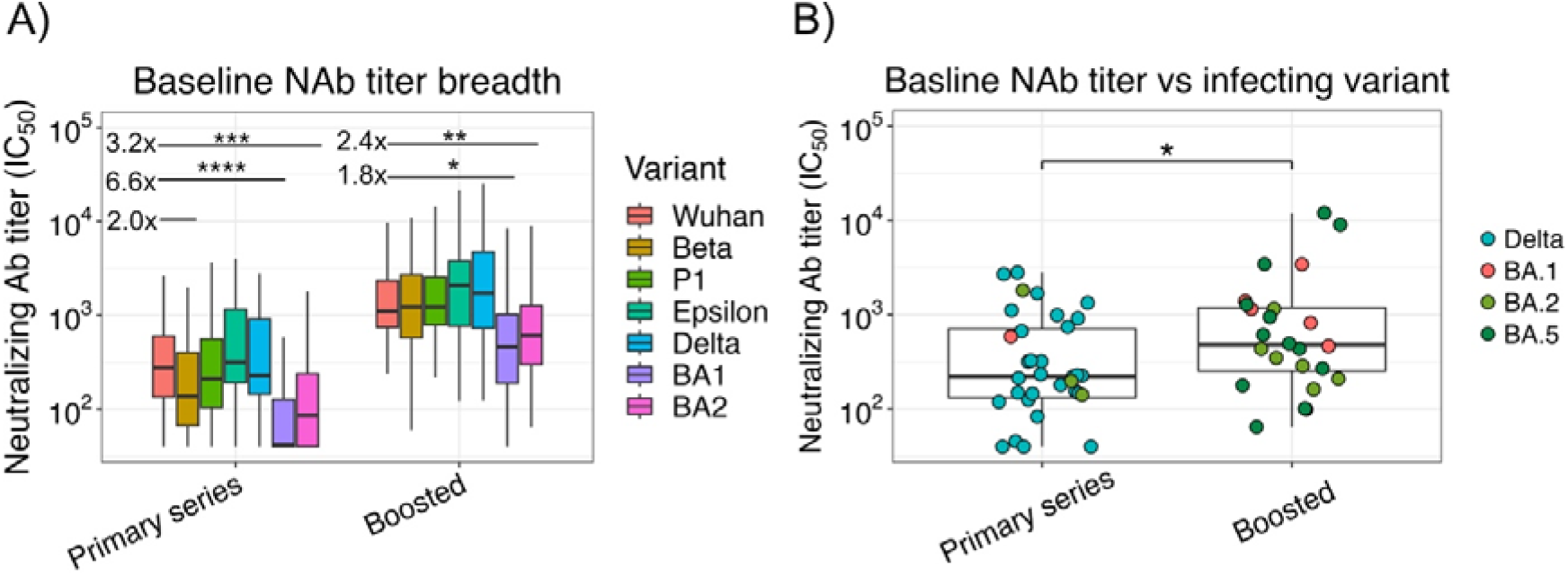
The magnitude and breadth of baseline NAb titers in participants with PVI. A) NAb titers targeting SARS-CoV-2 variants compared to Wuhan-Hu-1 in participants with PVI who received a primary vaccine series or monovalent booster vaccinations. N=22 participants with recruitment specimens taken >7 PSO were excluded from the analysis. B) Comparison of baseline NAb titers targeting the infecting variant (except for BA.5 infections for which responses against BA.2 are shown) between participants with PVIs who received a primary vaccine series or monovalent booster vaccinations. Statistical comparisons were made using a two-sided Wilcoxon rank sum test. Comparisons with a P value < 0.1 are shown. *P < 0.05, ** P < 0.01, ***P < 0.001 and ****P < 0.0005.

### Viral replication and duration of infection following PVIs are associated with baseline NAb titers in a variant-specific manner

Given that the strength of the baseline NAb titers differed by targeted variant, we next assessed the correlation between baseline NAb titers against the infecting variant (or BA.2 in the case of BA.5 infections) and features of viral replication dynamics (**Fig. 3**). In participants infected with Delta variants, we observed a significant negative correlation between baseline Delta-specific NAb titers and both maximum viral RNA load (*R*=-0.55, P<0.0069; **Fig. 3A**) and the duration of infectious virus shedding (R=-0.5, P=0.014; **Fig. 3B**). In participants infected with Omicron variants, baseline titers against the infecting variant were not associated with viral load or duration of infectious viral shedding (**Fig. 3C &D**).

**Figure 3.**
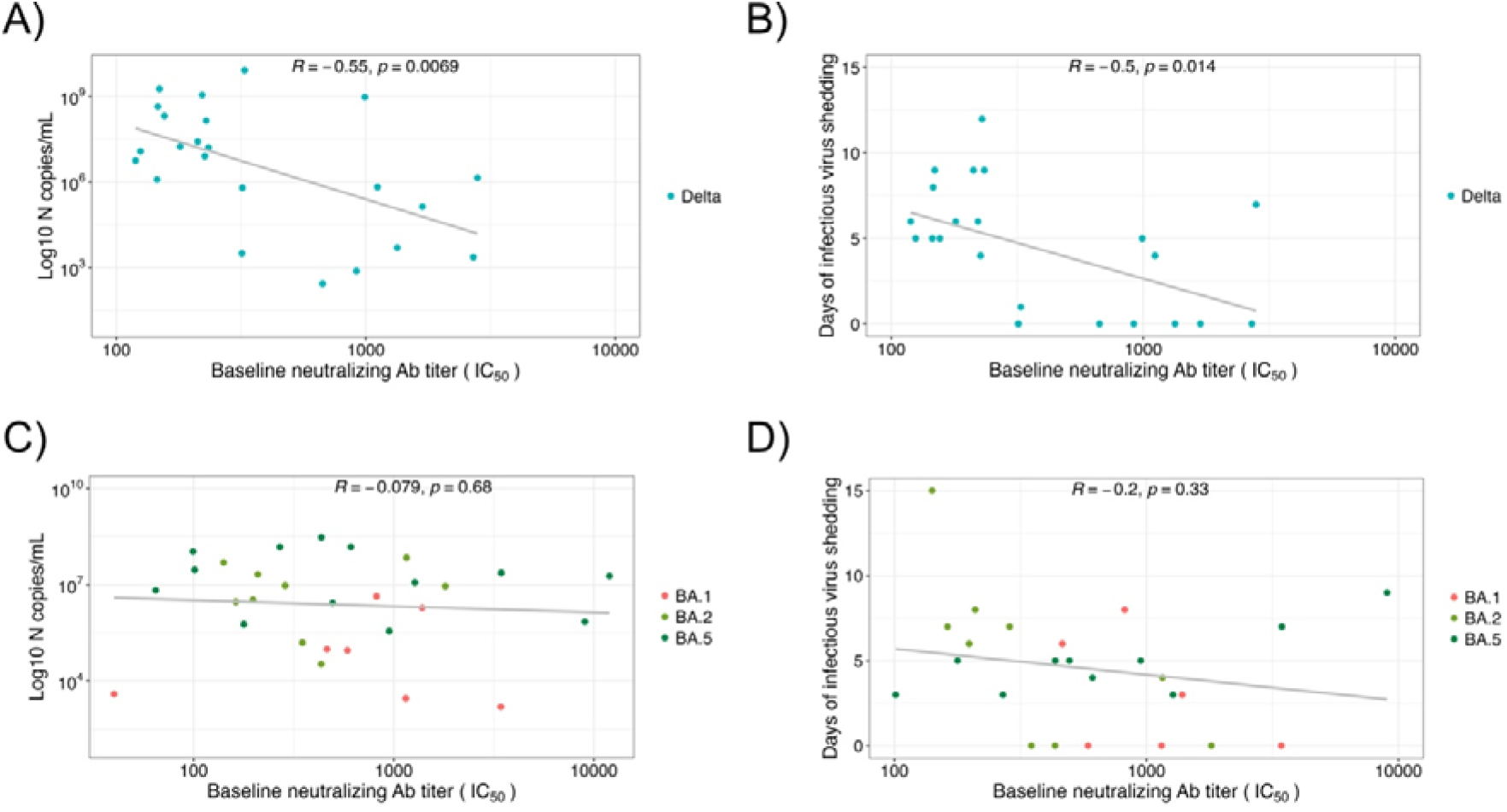
The relationship between baseline NAb titers and viral shedding outcomes. The correlation between baseline NAb titer targeting the infecting variant in vaccinated participants (N=29) infected with Delta variants and maximum RNA viral copies (A) and infectious virus shedding (B) over the study period. The correlation between baseline NAb titer targeting the infecting variant (except for participants with BA.5 infections for which responses were against BA.2) in vaccinated participants (N=29) infected with Omicron variants and maximum RNA viral copies (C) and infectious virus shedding (D) over the study period. Participants (N=9) with recruitment specimens taken on day 7 PSO or later were excluded. Pearson’s correlation coefficients and infecting variant are shown.

To further quantify the effect of baseline NAbs on virological outcomes following PVI (maximum viral load and duration of infectious shedding), we used multivariable linear regression in separate models for participants with Delta and Omicron infections. We adjusted for age and time since last vaccination (in Delta and Omicron infections) and, additionally, for Omicron variant (BA.1 BA.2 or BA.5) in Omicron infections (**Table 1**). We did not adjust for vaccine status in our two models, as vaccination status was colinear with variant infection in our cohort. Higher baseline NAb titers were independently associated with lower peak viral load and shorter duration of infectious viral shedding in Delta infections; per log_10_ increase in baseline NAb titer, we observed a -2.43 reduction in log_10_ maximum viral load (95% CI: -3.76, -1.11; P=0.0009; **Table 1**), and a -2.79 day reduction in duration of infectious virus shedding (95% CI: -4.99, -0.60; P=0.02; **Table 1**). However, in Omicron infections, no significant associations were observed between baseline NAb titers and maximum viral RNA load or the duration of infectious virus shedding (**Table 1**). In line with univariate results in figure 1D BA.1 infection was independently associated with reduced maximum viral RNA titers compared to BA.2/4 (P=0.01) and BA.5 infections (P=0.004).

**Table 1:**
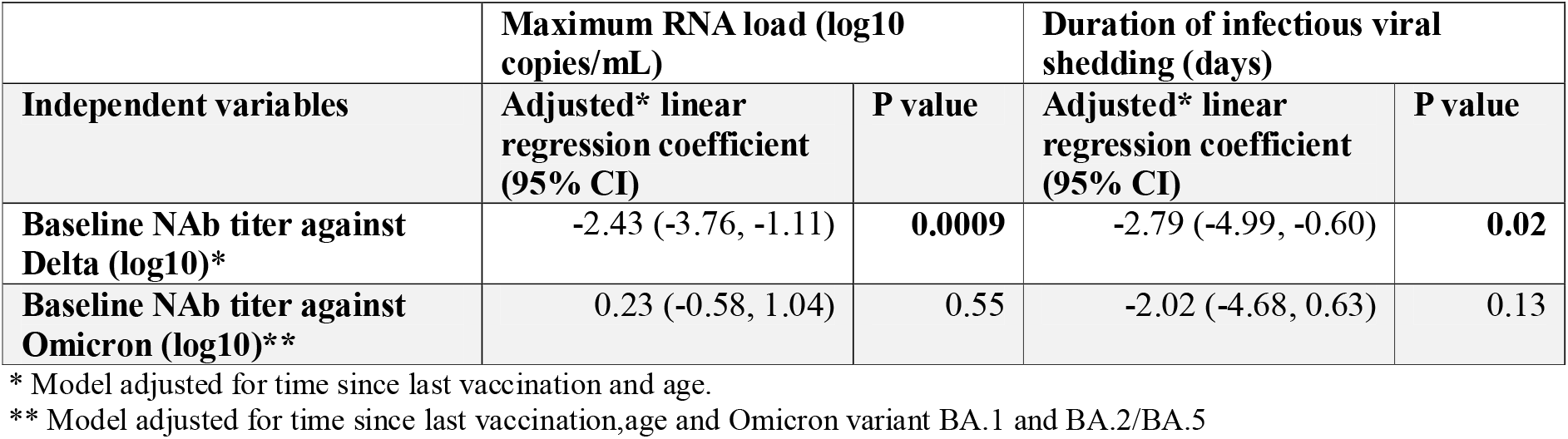
Effect of baseline neutralizing antibody titer on maximum viral RNA load and duration of infectious viral shedding in Delta and Omicron infections (N=29 participants in each group).

## DISCUSSION

Our findings show that following PVI (in persons vaccinated with ancestral spike antigens), higher baseline NAb titers are associated with accelerated viral clearance dynamics following infections with Delta variants, and we provide robust estimations to help quantify this effect. In addition, we find that in participants with PVI, baseline NAb titers targeting a range of variants up to BA.2 were increased in those who received booster vaccine doses compared to a primary series alone, as previously reported ^14,15^, and that booster vaccination was associated with a greater breadth of response including robust responses against early circulating immune escape variants Beta and P1. However, significantly reduced baseline NAb titers targeting BA.1 and BA.2 variants were observed in all vaccinated participants, and viral clearance was not influenced by NAb titers in participants infected with Omicron variants. This suggests that NAbs generated through first generation vaccines are limited in their ability to target conserved epitopes in the spike protein of Omicron variants and support the use of booster vaccination with updated antigens that may further broaden the NAb response.

To accurately estimate maximum RNA titers and duration of shedding of infectious virus we analyzed nasal specimens collected daily from participants. Studies assessing the effect of vaccination status on viral shedding dynamics have reported contrasting results, with initial studies suggesting an impact on peak RNA viral loads compared to unvaccinated individuals following infection with Alpha variants^16^. However, reports of Delta and Omicron BA.1 infections in outpatient cohorts^8,9,17^ and in individuals 2-6 months after receiving the last vaccine dose^7^, suggest no substantial impact on viral RNA load. Accordingly, our analysis including unvaccinated individuals, indicated no association between vaccination status and the virological outcomes assessed in our study. Taken together, this suggest that any positive impacts of first-generation vaccines on viral shedding outcomes and consequently, on transmission, may be rapidly lost (within 6 months following the last vaccine dose). In contrast to the effect of vaccination, infection with Omicron BA.1 was associated, in both univariate and multivariable analyses, with reduced maximum RNA titers compared to pre-Delta, Delta and BA.5 infection, as previously suggested^18^. The mechanisms by which BA.1 remained infectious at low viral loads^19^, achieving attack rates that exceeded those of other variants^20^ with lower peak viral titers remains to be fully understood, although there is evidence that mutations accumulated in Spike reduce the barrier to infection through accelerated early infection kinetics and evasion of innate immunity compared to Delta and pre-Delta variants^21,22^.

Our study has some limitations. We were not able to measure baseline NAb titers to BA.5 or to currently circulating Omicron variants, underscoring the importance of continued monitoring of the strength and breadth of immune responses against emergent antigenically divergent lineages such as JN.1. We used BA.2 as a proxy to estimate NAb responses to BA.5 in those who had BA.5 infections. Given that BA.5 descended from BA.2 and that all vaccinations contained ancestral spike antigens, we expect the immune escape phenotype of BA.5 to impact the results in figure 3C, 3D and Table 1 in similar manner to BA.2. In addition, we were not able evaluate the impact of vaccine boosters containing updated spike antigens, such as those containing sequences from XBB.1.5. Recent data suggest a further broadening of the NAb response following monovalent XBB.1.5 booster with robust targeting of JN.1^23^. Whether efficient broadening of baseline NAb titers by updated vaccine antigens leads to a restored correlation with viral shedding outcomes in PVI with currently circulating variants warrants investigation. Similarly, defining the antigenic distance^23,24^ between vaccine antigens and the infecting variant at which the correlation between baseline NAb titers and viral clearance dynamics is lost warrants future study.

The quantification of the effect of baseline NAbs on viral clearance may help parameterize mathematical models of transmission dynamics^25,26^ or immunobridging studies^27^ following the development of novel vaccines or the emergence of novel immune escape variants. To this end, further assessments of the relationship between circulating NAbs, mucosal NAbs and viral shedding dynamics are warranted, as the importance of the induction of mucosal immunity to limit SARS-CoV-2 transmission is increasingly recognized^28–30^. Lastly, recent studies have found early induction of memory T cell responses following PVI and have linked these responses with protective immunity^31^. Quantification of the added contribution of memory T cells to the early control of viral replication is critical to understanding the correlates of protection against SARS-CoV-2 infection.

## METHODS

### Study population and design

This observational longitudinal cohort, recruited in the San Francisco Bay area, was designed to characterize virological, immunological, and clinical outcomes of SARS-CoV-2 infection and household transmission dynamics, as previously detailed^11^. Briefly, both index cases and household contacts were recruited if an index case was identified from individuals with a positive health provider-ordered SARS-CoV-2 nucleic acid amplification test result on a nasopharyngeal or oropharyngeal (NP/OP) specimen done at UCSF-affiliated health facilities.. Index cases were defined as being infected with SARS-CoV-2 within 5 days of symptom onset by clinical nucleic acid amplification tests at study health facilities. Household contacts were defined as cohabitants of the index case that did not report COVID-19-like symptoms in the preceding week. Starting at enrollment, index cases and contacts self-collected nasal specimens daily for 2 weeks –with day 0 defined as the day of symptom onset of the index case– and on days 17, 19, 21 and 28. Specimens were stored at -20°C in a designated freezer provided to the participants, collected weekly by study staff, and stored at -80°C long-term. Venous blood specimens were collected at enrollment and during weekly home visits at days 9, 14, 21 and 28 post-symptom onset of the index case. The timing in days post symptom onset for each specimen was adjusted retrospectively for contact cases according to self-reporting. A survey was performed at or before enrollment to collect information on demographics, underlying conditions, prior infections, symptom start date and vaccine doses received. This activity was reviewed by UCSF and CDC, deemed not research, and was conducted consistent with applicable federal law and CDC policy (§See e.g., 45 C.F.R. part 46, 21 C.F.R. part 56; 42 U.S.C. §241(d); 5 U.S.C. §552a; 44 U.S.C. §3501 et seq)

### Neutralizing antibody response assay

The PhenoSense SARS CoV-2 nAb Assay (Monogram Biosciences, South San Francisco, CA, USA) was used to determine NAb titers as described previously^32,33^. Briefly, the assay was done using HIV-1 pseudotype virions expressing SARS-CoV-2 spike proteins from Wuhan-Hu-1, Beta, P1, Epsilon, Delta, BA.1 and BA.2. Virions were generated in HEK293 cells following co-transfection of a spike-encoding vector with an HIV-1 genomic vector expressing firefly luciferase. Reduction is luciferase activity in infected HEK293 cells expressing human Ace2 and TMPRSS2, following preincubation of pseudoviriones with serial solutions of patient plasma, was used to determine the 50% infectious dose (ID50). NAb titers were determined at all available timepoints (days 7, 14, 21 and 28). Maximum NAb titer was defined as the NAb titer on the day with the highest NAb titers against the variant of interest, for each participant.

### RNA extraction

RNA extraction from 200uL of nasal specimens was done using the KingFisher (Thermo Scientific) automated extraction instrument and the MagMAX Viral/Pathogen Nucleic Acid Isolation Kit (Thermo Scientific) following the manufacturer’s instructions as previously described^8^. For confirmatory RT-qPCR the Quick-DNA/RNA Viral MagBead kit (Zymo) was used as previously described^8^.

### RT-qPCR assay

For each RT-qPCR reaction, 4μL of RNA sample were mixed with 5μL 2x Luna Universal Probe One-Step Reaction Mix, 0.5μL 20x WarmStart RT Enzyme Mix (NEB), 0.5μL of target gene specific forward and reverse primers and probe mix as previously described^8^. RT-qPCR were run for SARS-CoV2 N and E and for host mRNA RNaseP as a control for RNA extraction. 8μM each of forward and reverse primers and 4μM probe for E; 5.6μM each of forward and reverse primers and 1.4μM probe for N; and 4μM each of forward and reverse primers and 1μM probe for RNaseP were used per reaction. Each 96 well RT-qPCR plate was run with a 10-fold serial dilution of an equal mix of plasmids containing a full copy of nucleocapsid (N) and envelope (E) genes (IDT), as an absolute standard for the calculation of RNA copies and primer efficiency assessment. RTqPCR were run on a CFX Connect Real-Time PCR detection system (Biorad) with the following settings: 55 □C for 10 min, 95 □C for 1min, and then cycled 40 times at 95 □C for 10s followed by 60 □C for 30s. Probe fluorescence was measured at the end of each cycle. All probes, primers and standards were purchased from IDT. A sample was considered to contain SARS-CoV-2 RNA if both N and E were detected at Ct <40. To control for the quality of self-sampling, RNAse P Ct values 2 standard deviations from the mean of all samples were repeated or excluded. Maximum RNA viral load was defined as the RNA value on the day with the highest RNA viral load for each participant.

### Cytopathic effect (CPE) assay

All anterior nares samples up to 14 days post symptom onset (PSO) of the index case were assayed for their ability to induce a cytopathic effect (CPE). In cases where CPE was still positive within days 11–14, we continued to test samples beyond day 14 until three consecutive samples were CPE negative. As described previously^8^, CPE was assessed on Vero-hACE2-TMPRSS2 cells. Cells were maintained at 37 □C and 5% CO2 in Dulbecco’s Modified Eagle medium (DMEM; Gibco) supplemented with 10% fetal calf serum, 100ug/mL penicillin and streptomycin (Gibco) and 10μg/mL of puromycin (Gibco). 200uL of nasal specimen was added to a well of a 96-well plate and serially diluted 1:1 with DMEM supplemented with 1x penicillin/streptomycin over two additional wells. 100uL of freshly trypsinized cells, resuspended in infection media (made as above but with 2x penicillin/streptomycin, 5ug/ mL amphotericin B [Bioworld] and no puromycin) at 2.5 x 10^5^ cells/mL, were added to each sample dilution. Cells were cultured at 37 □C and 5% CO2 and checked for CPE from day 2 to 5. After 5 days of incubation, the supernatant (200uL) from one well from each dilution series was mixed 1:1 with 2x RNA/DNA Shield (Zymo) for viral inactivation and RNA extraction as described above. Among specimens with visible CPE, the presence of infectious SARS-CoV-2 was confirmed by RT-qPCR using N primers as described above. Duration of infectious viral shedding was defined as days between symptom onset and the last day of CPE positivity for each participant. All assays were done in the BSL3 facility at Genentech Hall, UCSF, following the study protocol that had received Biosafety Use Authorization.

### Sequencing

The ARTIC Network amplicon-based sequencing protocol for SARS-CoV-2 was used to sequence the nasal specimen with the highest copies of viral RNA per participant. Thawed RNA specimens were converted to cDNA using the Luna RT mix (NEB). Arctic multiplex PCR primer pools (IDT) (versions 4.1 and 5.3.2) were used to generate amplicons that were barcoded using the Native Barcode expansion kits 1–24 (Nanopore), pooled and used for adaptor ligation. Libraries were run on a MinION sequencer (Oxford Nanopore Technologies) for 16 hours. Consensus sequences were generated using the nCoV-2019 novel coronavirus bioinformatics protocol using the MinIon Pipeline. Lineage determination was done using the online Pangolin COVID-19 Lineage Assigner.

### Statistical analysis

Categorical data was summarized as frequencies of the total population. Continuous data was summarized with median values and interquartile ranges. Comparisons of group medians was done using two-sided unpaired Wilcox rank sum tests. We assessed the relationship between NAb responses and viral shedding dynamics, stratified by variant and vaccination status, in unadjusted and adjusted analyses. Stratified by Delta vs Omicron infections, we used multivariable linear regression to assess the effect of baseline NAbs on viral outcomes (maximum RNA load, continuous variable; duration of infectious viral shedding, continuous), adjusting for age, time since vaccination (months), and Omicron sub-variant. Data was analyzed with custom scripts using R in RStudio (version2023.06.1+524).

## ACKNOWLEDGEMENTS

We would like to thank the participants for their time and efforts to make this study possible. The findings and conclusions in this report are those of the authors and do not necessarily represent the official position of the Centers for Disease Control and Prevention.

---

**Supp. table 1.** Participant characteristics (N=174).

| **Characteristic** | **SARS-CoV-2-Infected** | **SARS-CoV-2-Uninfected** |
| --- | --- | --- |
| **Total** | N=125 | N=49 |
| **Case** |  |  |
| Index case*, n (%) | 78 (62.4) | 0 (0.0) |
| Household contact (%) | 47 (37.6) | 49 (100.0) |
| **Sex, n (%)** |  |  |
| Female | 65 (52.0) | 25 (51.0) |
| Male | 60 (48.0) | 24 (49.0) |
| **Age category, n (%)** |  |  |
| 13-17 | 3 (2.4) | 3 (6.1) |
| 18-29 | 27 (21.6) | 11 (22.4) |
| 30-39 | 43 (34.4) | 13 (26.5) |
| 40-49 | 28 (22.4) | 12 (24.5) |
| 50-59 | 11 (8.8) | 7 (14.3) |
| >60 | 13 (10.4) | 3 (6.1) |
| **Race/Ethnicity, n (%)** |  |  |
| Hispanic or Latino Ethnicity | 28 (22.4) | 10 (20.4) |
| White | 54 (43.2) | 23 (46.9) |
| Black or African American | 4 (3.2) | 3 (6.1) |
| Asian | 18 (14.4) | 6 (12.2) |
| Pacific Islander/ Hawaiian | 2 (1.6) | 2 (4.1) |
| American Indian / Alaska Native | 2 (1.6) | 0 (0.0) |
| NA | 17 (13.6) | 5 (10.2) |
| **BMI, n (%)** |  |  |
| <25 | 51 (40.8) | 21 (43.0) |
| 25 to 29.9 | 32 (25.6) | 16 (32.7) |
| >30 | 25 (20.0) | 6 (12.2) |
| NA | 17 (13.6) | 6 (12.2) |
| **COVID-19 vaccination status\*\*, n (%)** |  |  |
| *Unvaccinated* | *57 (45.6)* | *21 (42.9)* |
| *Primary series* | *37 (29.6)* | *12 (24.5)* |
| Median (IQR) days since most recent dose | 182 (123-229) | 115 (89-150) |
| Most recent dose <180 days before baseline | 23 (62.2) | 12 (100) |
| Most recent dose 180+ days before baseline | 14 (37.8) | 0 (0) |
| *One or two boosters* | *31 (24.8)* | *15 (30.6)* |
| Median (IQR) days since most recent dose | 178 (107-227) | 109 (43-191) |
| Most recent dose <180 days before baseline | 13 (41.9) | 10 (66.7) |
| Most recent dose 180+ days before baseline | 18 (58.1) | 5 (33.3) |

| **Variant, n (%)\*\*\*** |  |  |
| --- | --- | --- |
| Pre-VOC | 45 (36.0) | NA |
| Epsilon | 4 (3.2) | NA |
| Delta | 34 (27.2) | NA |
| Omicron BA.1 | 10 (8.0) | NA |
| Omicron BA.2, BA.4 or BA.5 | 32 (25.6) | NA |

\*SARS-CoV-2 infected participant approached in each household
\*\* Last vaccine dose 4 weeks or greater before study enrolment. 170/174 participants received either BNT162b2 (Pfizer) or mRNA-1273 (Moderna) primary vaccine series and when indicated, monovalent boosters. Two infected individuals received a Ad26.COV2.S (Janssen) primary series, one uninfected participant received a Ad26.COV2.S (Janssen) primary series and one uninfected participant a AZD1222 (AstraZeneca) primary series.
\*\*\* Determined by sequencing of SARS-CoV-2 RNA from the participant or cohabiting participant's nasal specimen.

**Supplementary figure 1.**
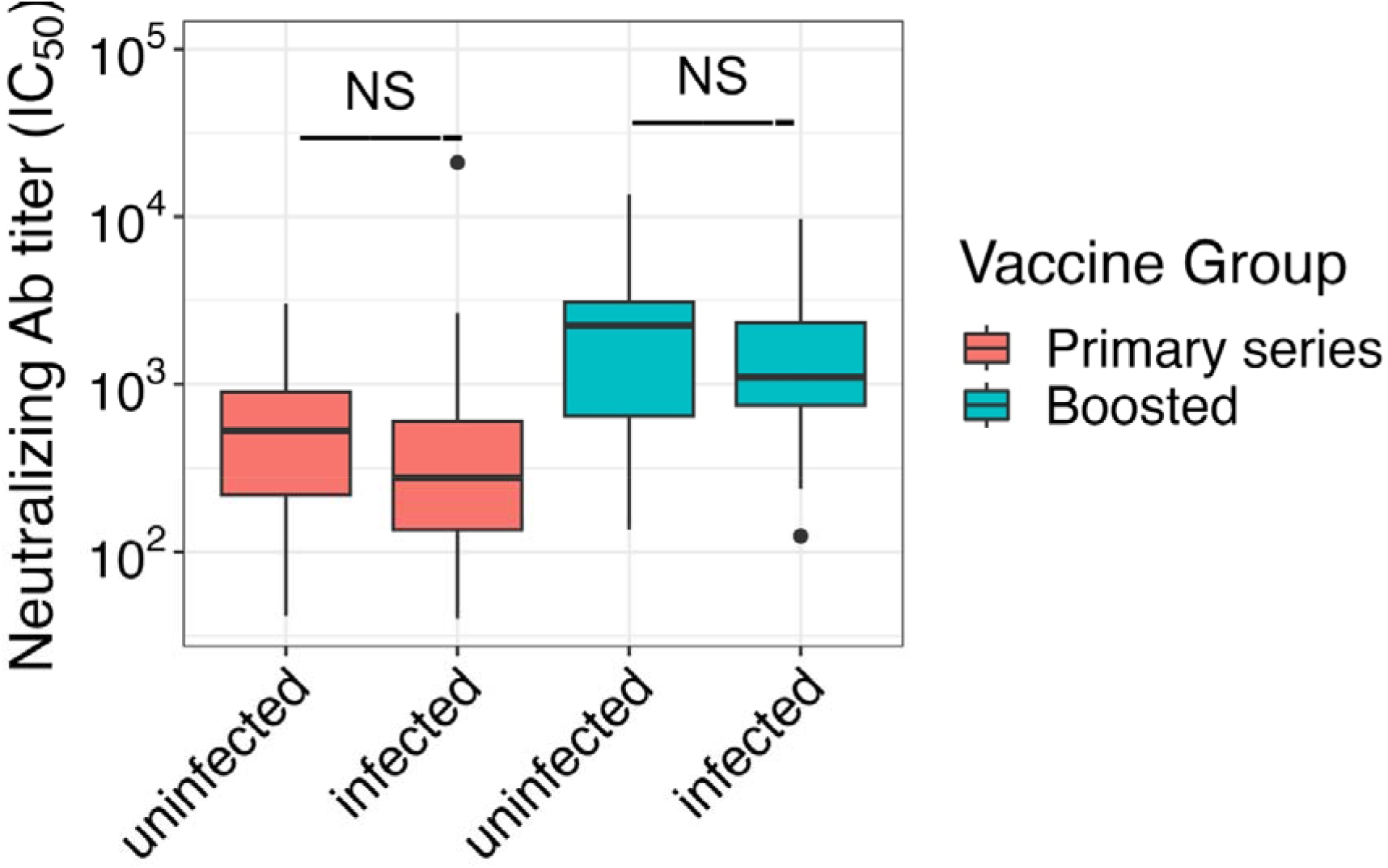
NAb titers collected at recruitment in uninfected and infected participants. Comparison of NAb titers in recruitment specimens against Wuhan-Hu-1 in infected and uninfected participants stratified by vaccination status. Recruitment specimens were collected <7 days following symptom onset in infected individuals. Statistical comparisons were done within vaccination groups by Wilcox test. NS, not significant.

**Supplementary figure 2.**
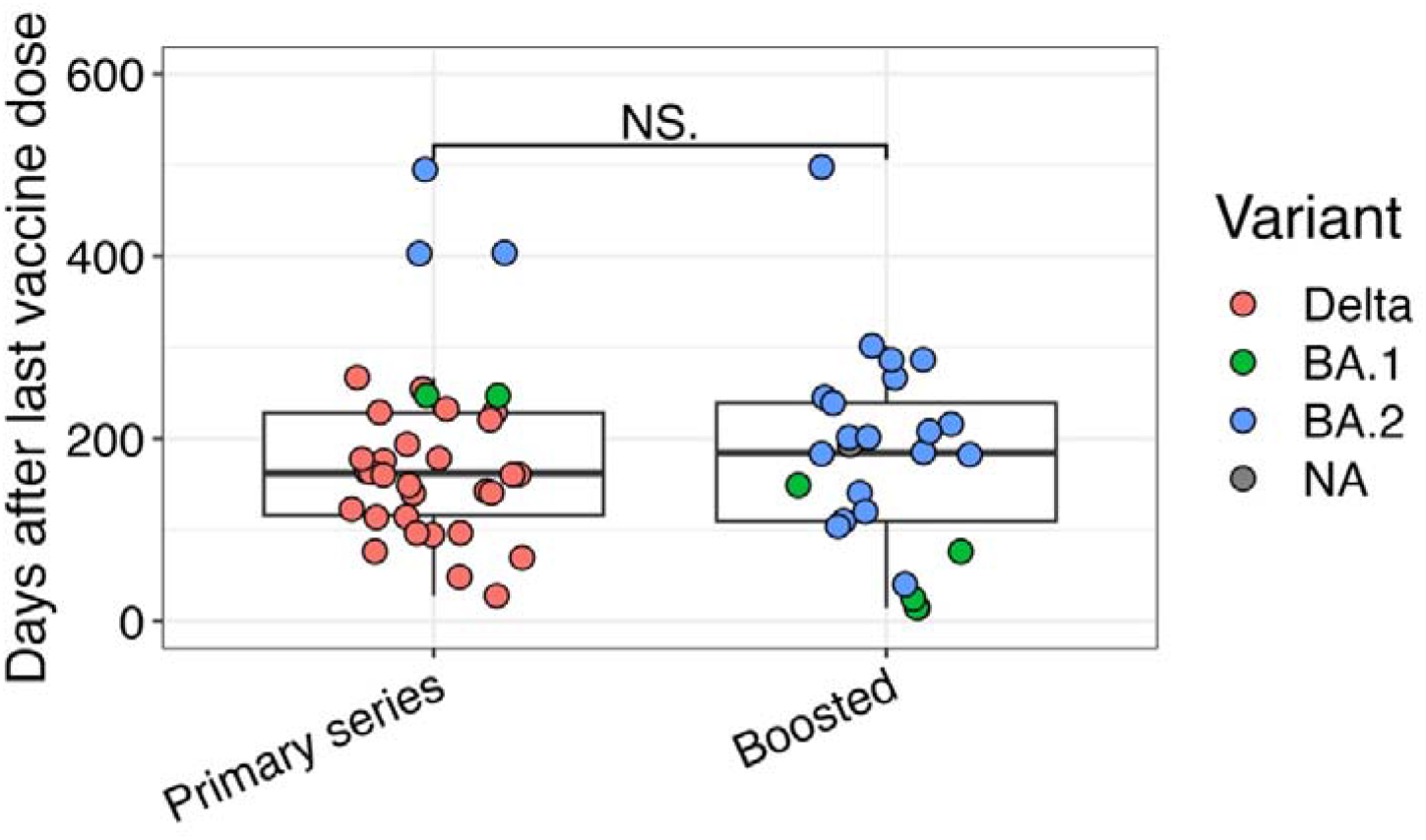
Time since last vaccine dose in vaccinated participants. Comparison of the days since the last vaccine dose in participants who received a primary vaccine series or booster vaccinations. Statistical comparison done by Wilcox test. NS, not significant.

